# Stress granules promote chemoresistance by triggering cellular quiescence

**DOI:** 10.1101/2022.02.22.481503

**Authors:** Anthony Khong, Nina Ripin, Luisa Macedo de Vasconcelos, Sabrina Spencer, Roy Parker

## Abstract

Cells respond to cellular stress by forming stress granules, molecular condensates containing non-translating messenger ribonucleoproteins. Stress granules form during chemotherapy and promote cell survival and chemoresistance, although the mechanism of this effect is not understood. We provide several lines of evidence that stress granules enhance cell survival by promoting cellular quiescence. First, we see a correlation between spontaneous stress granule formation and cell-cycle exit under non-stress conditions. Second, cells deficient in proteins required for stress granule formation (G3BP1/2) are less likely to exit the cell cycle under non-stress, stress, and chemotherapeutic conditions. Third, rescuing stress granule formation in G3BP1/2 knockout cells restores the fraction of cycling cells to wild-type levels. Finally, cells with enhanced stress granule formation (ddx6 knockout cells) show an increased propensity to exit the cell cycle. These results suggest that stress granules are important regulators of cellular quiescence, which could enable the identification of new anti-chemoresistance therapies that target stress granules.

## INTRODUCTION

Stress granules are large molecular condensates that form under a variety of cellular stress conditions including oxidative stress, heat shock, osmotic stress, and chemotherapeutics (Anderson et al., 2015; Protter and Parker, 2016). During cellular stress, the integrated stress response pathway is activated, which triggers eIF2α phosphorylation resulting in global translational initiation repression. Subsequently, most mRNAs are released from translating ribosomes resulting in the accumulation of non-translating messenger ribonucleoproteins (mRNPs), which becomes the core assemblers and constituents of stress granules (Jain et al., 2016; Khong et al., 2017; Markmiller et al., 2018; Namkoong et al., 2018; Youn et al., 2018).

Numerous studies indicate an intimate link between stress granules and cancer biology (Anderson *et al*., 2015; Lavalée et al., 2021; Song and Grabocka, 2020). First, stress granules are found in a variety of tumors (Grabocka and Bar-Sagi, 2016; Somasekharan et al., 2015; Valentin-Vega et al., 2016). Second, pro-tumorigenic signaling pathways promote stress granule assembly (Song and Grabocka, 2020). Third, poor patient prognosis correlates with the high expression levels of the many core stress granule assembly proteins (Aucagne et al., 2017; Grabocka and Bar-Sagi, 2016; Somasekharan *et al*., 2015). Fourth, many chemotherapeutic agents trigger stress granule formation (Adjibade et al., 2015; Bittencourt et al., 2019; Fournier et al., 2010; Grabocka and Bar-Sagi, 2016; Kaehler et al., 2014; Lin et al., 2019; Szaflarski et al., 2016; Zhao et al., 2021). However, how stress granules contribute to cancer biology is not well understood. One of the prevailing models is that stress granules help cancer cells adapt to a variety of cellular stress elicited by the tumor microenvironment and promote resistance to chemotherapeutic drugs (Anderson *et al*., 2015; Lavalée *et al*., 2021; Song and Grabocka, 2020).

Several mechanistic molecular models have been proposed to explain how stress granules may contribute to cell survival during stress. One common theme in many of these models is the sequestration of pro-apoptotic proteins to stress granules which leads to the dampening of apoptotic signaling pathways. This includes the pro-apoptotic proteins RACK1 (Arimoto et al., 2008), TRAF2 (Kim et al., 2005), RhoA-Rock1 complex (Tsai and Wei, 2010), YWHAX (Zhao *et al*., 2021), BAX (Zhao *et al*., 2021), and mTORC1 constituents (Thedieck et al., 2013; Wippich et al., 2013). Another common theme is that stress granules may also promote cell survival by reducing reactive oxygen species in cells through a variety of mechanisms (Amen and Kaganovich, 2021; Takahashi et al., 2013).

In this manuscript, we describe a new mechanism by which stress granules promote cell survival and chemoresistance by promoting cellular quiescence. Cellular quiescence is a state of reversible growth arrest with reduced transcription and translation from which cells can re-enter the proliferative state. Cellular quiescence is thought to promote cell survival by protecting damage through reducing reactive oxygen species by favoring glycolysis over oxidative phosphorylation, upregulating antioxidant genes, increasing autophagic flux, and downregulating apoptosis genes (Coller et al., 2006; Marescal and Cheeseman, 2020). Moreover, cellular quiescence also protects cells against chemotherapeutics, which in general target cycling cells.

A recently published article isolated and described a subpopulation of MCF10A cells that are slow-cycling (Min and Spencer, 2019). This subpopulation of cells frequently passes through a transient quiescent (G0) state even in untreated conditions. This state is marked by low Cyclin-Dependent Kinase 2 activity, high p21 levels, and hypo-phosphorylated Rb and is triggered by activation of stress-response pathways including p53 activation, or eIF2α phosphorylation and the inhibition of translation. Since inhibition of translation should lead to the assembly of stress granules, we investigated if stress granules are involved in establishing or maintaining quiescence.

We discovered that stress granules are involved in promoting or maintaining cell-cycle exit (quiescence) based on multiple observations. First, untreated cells containing spontaneous stress granules are more likely to be quiescent than cells without stress granules. Second, cells deficient in core stress granule proteins (G3BP1/2 double knockout) have a reduced fraction of quiescent cells under non-stress and stress conditions. Third, the number of G3BP1/2 dKO cells in quiescence can be rescued by sorbitol or high levels of Hippuristanol, manipulations that rescue stress granule formation in the absence of G3BP1/2 proteins (Kedersha et al., 2016; Tauber et al., 2020). Fourth, cells deficient in DDX6 have increased stress granule formation, which then leads to greater levels of cellular quiescence. Fifth, the chemotherapeutic Vinorelbine induces less quiescence in G3BP1/2 dKO compared with wild-type cells. These observations argue that stress granules promote entry into quiescence, which then leads to cell survival and chemoresistance.

## RESULTS

### Cells lacking G3BP1/2 genes show reduced quiescent cells

Given the relationship between the integrated stress response and quiescence (Min and Spencer, 2019), we first asked if the presence of stress granules is associated with spontaneous quiescence. To examine stress granule assembly, we used a G3BP1/2 double knock out (dKO) U-2 OS cell line (Kedersha *et al*., 2016), as well as cells which have been rescued with a GFP-G3BP1 transgene by lentivirus transduction (GFP-G3BP1 + G3BP1/2 dKO U-2 OS cells). This transgene is expressed at comparable levels to the G3BP1 proteins found in wildtype U-2 OS cells (Figure 1A). We then stained cells with an antibody that recognizes Rb phosphorylation. Rb is hypo-phosphorylated in quiescent cells and becomes hyper-phosphorylated once cells cross the Restriction Point and commit to the cell cycle (Weinberg, 1995). Since spontaneous stress granule formation is expected to be a low-frequency event, we used a high-content screening microscope to image thousands of cells rapidly.

**Figure 1.**
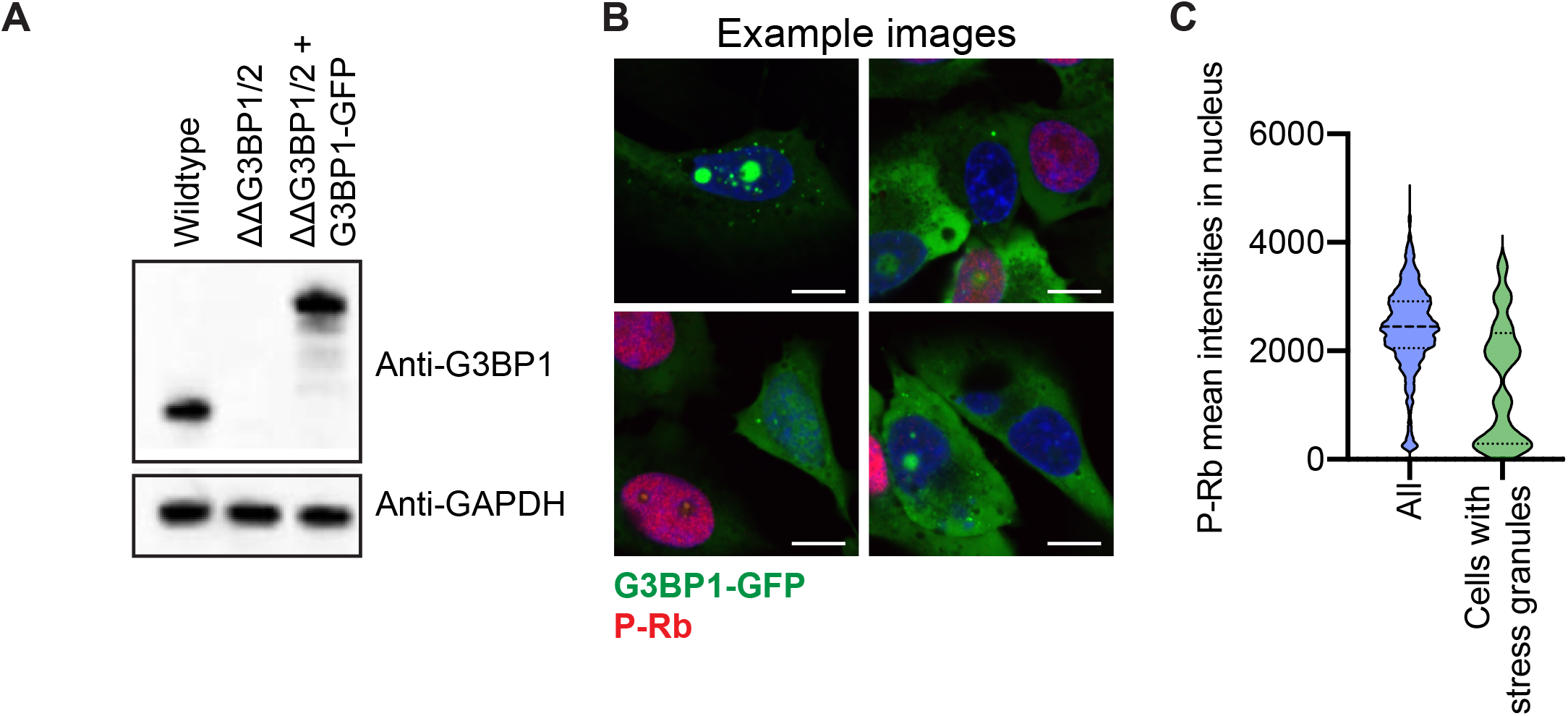
Stress granule formation correlates with low phospho-Rb levels. **(A)** G3BP1 and GAPDH immunoblots from wildtype, G3BP1/2 dKO U-2OS, and GFP-G3BP1 + G3BP1/2 dKO U-2 OS. Scale bar = 10 µm. **(B)** Images of cells forming spontaneous stress granules (GFP-G3BP1 puncta) in non-stress conditions (Red = phospho-Rb, Green= GFP-G3BP1, Blue = DAPI). **(C)** Violin plot distribution of phospho-Rb in all cells and cells with visible stress granules. The total number of cells is 1947. Cells containing stress granules are 20.

By comparing cells that form stress granules and the phosphorylation status of Rb, we discovered cells with spontaneous stress granules have reduced Rb phosphorylation (Figure 1C). These spontaneous stress granules are observed in approximately 1% in GFP-G3BP1 + G3BP1/2 dKO U-2 OS cells (Figure 1B) and are typically smaller than arsenite-induced stress granules (Figure 2A). These results suggest that stress granules might be involved in establishing or maintaining cellular quiescence.

**Figure 2.**
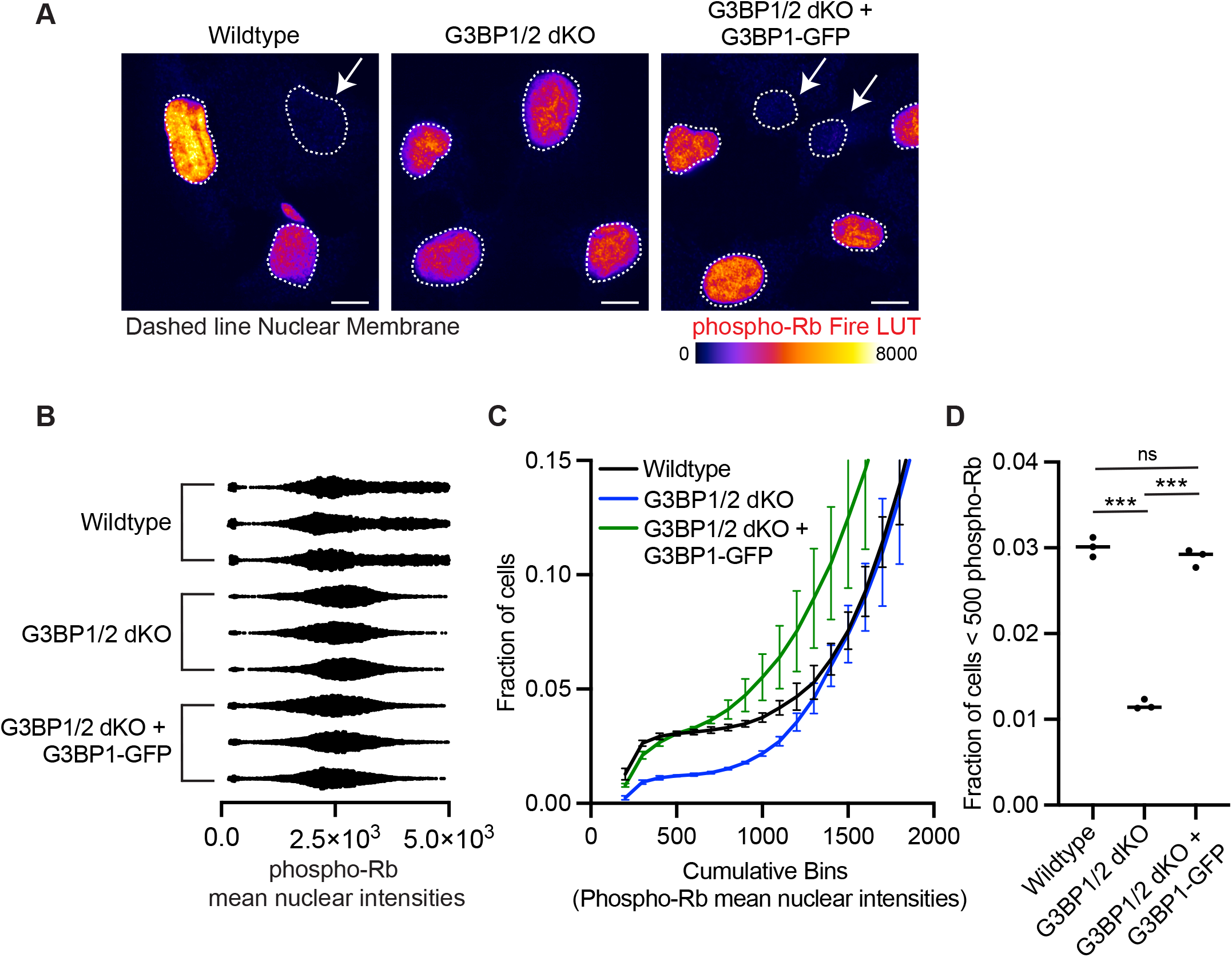
G3BP1/2 genes promote a greater fraction of cells with low phospho-Rb levels. **(A)** Representative immunofluorescence phospho-Rb images of wildtype, G3BP1/2 dKO U-2 OS, and GFP-G3BP1 + G3BP1/2 dKO U-2 OS cells. The phospho-Rb immunofluorescence is converted to Fire LUT and the dashed lines indicate the nuclear membrane. Arrowheads denote quiescent cells. Scale bar = 10 µm. **(B)** Mean phospho-Rb nuclear intensity scatterplot of wildtype, G3BP1/2 dKO U-2OS, and GFP-G3BP1 + G3BP1/2 dKO U-2 OS cells. Each dot represents a biological replicate. At least 3487 cells were counted for each biological replicate. **(C)** Cumulative frequency plot from 0 to 2000 phospho-Rb mean nuclear intensities from Figure 2B. **(D)** Scatterplot of the fraction of cells with phospho-Rb mean nuclear intensity less than 500 from Figure 2B. Three biological replicates were performed. ns = P-value > 0.05; *** = P-value < 0.001.

We next examined if stress granules are involved in promoting or maintaining the quiescent phase by examining cells deficient in G3BP1/2 proteins. G3BP1/2 proteins are critical for stress granule formation and knockout of G3BP1/2 genes severely limits stress granule formation under a variety of stress conditions and would be expected to affect these spontaneously forming stress granules (Kedersha *et al*., 2016). We, therefore, quantified the phospho-Rb signal in wildtype, G3BP1/2 dKO, and G3BP1/2 dKO + GFP-G3BP1 U-2 OS cells (Figure 2A-B). A key observation is that wildtype cells have a greater fraction of cells with hypo-phosphorylated-Rb than G3BP1/2 dKO U-2 OS cells (Figure 2B). While both wildtype and G3BP1/2 dKO U-2 OS cells show a wide distribution of phospho-Rb levels (Figure 2B), the G3BP1/2 dKO U-2 OS cells lack the population of cells containing hypo-phosphorylated Rb. This difference can also be observed using a cumulative frequency plot of phospho-Rb intensities (Figure 2C), or a strict threshold scatterplot for phospho-Rb (Figure 2D). Importantly, the introduction of the GFP-G3BP1 transgene into the G3BP1/2 dKO U-2 OS cells restores the fraction of cells with hypo-phospho-Rb to a level comparable to wildtype U-2 OS cells (Figure 2B-D). These results indicate the G3BP1/2 genes are involved in promoting and/or maintaining quiescence.

We next asked if stress granules are also involved in promoting and/or maintaining the quiescent phase of the cell cycle under arsenite conditions. Arsenite is known to robustly inhibit translation and induce stress granule formation (Kedersha et al., 1999). We, therefore, treated wildtype, G3BP1/2 dKO, and GFP-G3BP1 + G3BP1/2 dKO U-2 OS cells with 100 µM arsenite for 6 hours. This condition induces stress granules robustly after 1h treatment in wildtype and GFP-G3BP1 + G3BP1/2 dKO U-2 OS cells, but not in G3BP1/2 dKO U-2 OS cells (Figure 3A-B). If stress granules promote quiescence, one predicts that arsenite treatment should lead to an increase in hypo-phosphorylated Rb cells in wild-type cells, and that population should be reduced in the dKO G3BP1/2 U-2 OS cells. Indeed, we observed a greater percentage of cells in quiescence in wildtype and GFP-G3BP1 + dKO G3BP1/2 U-2 OS cells compared to dKO G3BP1/2 U-2 OS cells (Figure 3C-F) after 6h of 100µM arsenite treatment. These results indicate the G3BP1/2 proteins are also involved in promoting or maintaining quiescence under stress granule-inducing arsenite conditions.

**Figure 3.**
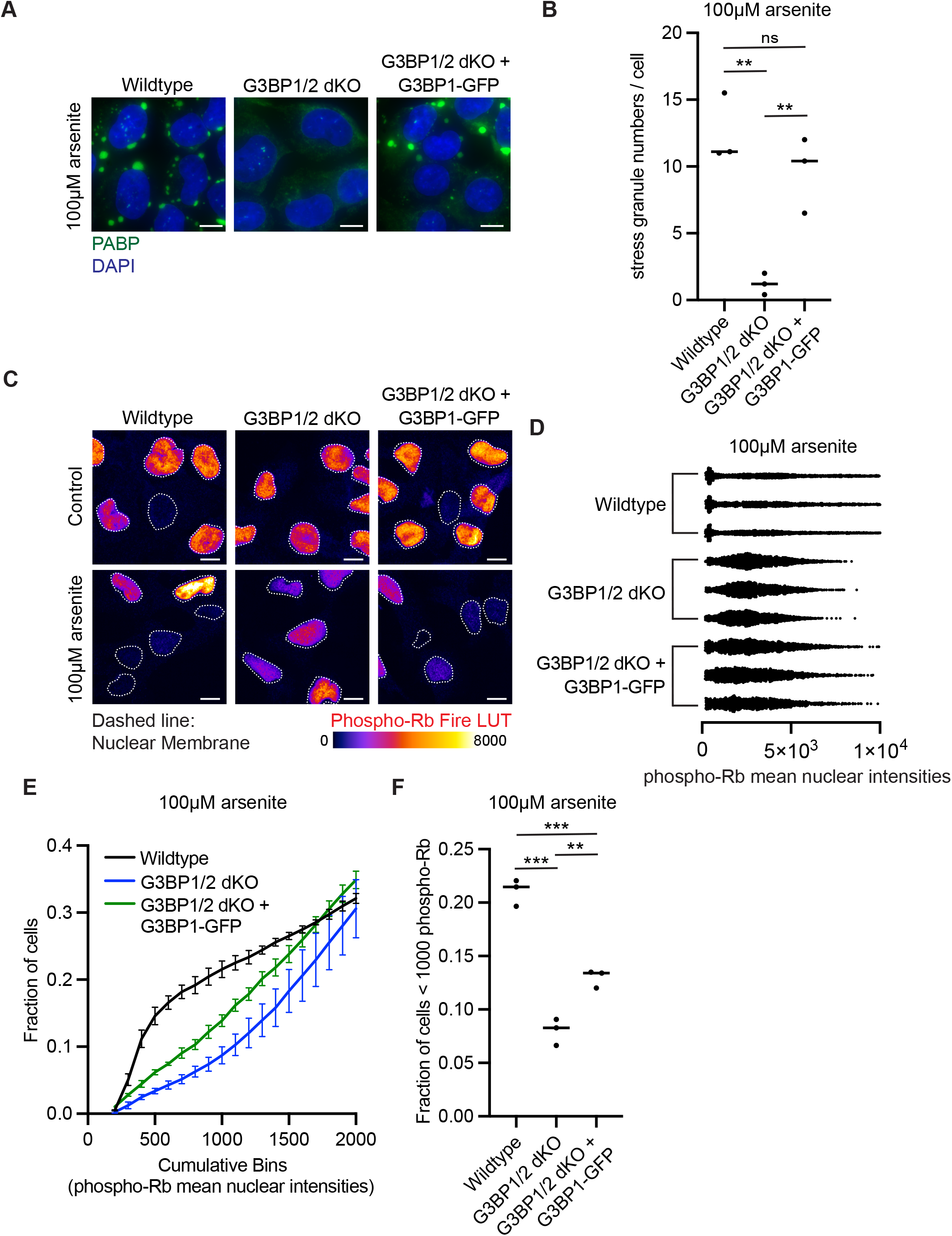
G3BP1/2 genes promote a greater fraction of cells with low phospho-Rb levels when treated with 100 µM arsenite. **(A)** Representative PABP immunofluorescence images of wildtype, G3BP1/2 dKO, and GFP-G3BP1 + G3BP1/2 dKO U-2 OS cells treated with 100 µM arsenite for 1 hour. Scale bar = 5 µm. **(B)** Scatterplot of the number of stress granules per cell from wildtype, G3BP1/2 dKO, and GFP-G3BP1 + G3BP1/2 dKO U-2 OS cells treated with 100 µM arsenite for 1 hour. Three biological replicates were performed. 10 cells were counted for each biological replicate. ns = not significant; ** = P-value < 0.001. **(C)** Representative phospho-Rb immunofluorescence images of wildtype, G3BP1/2 dKO, and GFP-G3BP1 + G3BP1/2 dKO U-2 OS cells treated with 100 µM arsenite for 6 hours or left untreated. The phospho-Rb immunofluorescence is converted to Fire LUT and the dashed lines indicate the nuclear membrane. Scale bar is 10 µm. **(D)** Mean phospho-Rb nuclear intensity scatterplot of wildtype, G3BP1/2 dKO U-2OS, and GFP-G3BP1 + G3BP1/2 dKO U-2 OS cells. Each dot represents a biological replicate. At least 1418 cells were counted for each biological replicate. **(E)** Cumulative frequency plot of phospho-Rb mean nuclear intensities from 0 to 2000 phospho-Rb mean nuclear intensities from Figure 3D. Error bars represent the three biological replicates from Figure 3D. **(F)** Scatterplot of the fraction of cells with phospho-Rb mean nuclear intensity less than 1000 from Figure 3D. Three biological replicates were performed. ** = P-value < 0.01; *** = P-value < 0.001.

### G3BP1/2 proteins promote quiescent cells through stress granule formation

In principle, G3BP1/2 proteins could promote quiescence by triggering stress granule formation, or by a G3BP1/2 function independent of their role in stress granule formation (Laver et al., 2020; Meyer et al., 2020; Prentzell et al., 2021). To distinguish if G3BP1/2 proteins affect quiescence through stress granules or independently, we examined how sorbitol affected the formation of quiescent cells. Sorbitol triggers stress granule formation independent of G3BP1/2 proteins (Figure 4A-B) (Kedersha *et al*., 2016; Matheny et al., 2021). We stressed wildtype and G3BP1/2 dKO U-2 OS for 6 hours with 0.5 M sorbitol concentrations and stained for phospho-Rb (Figure 4C). If the role of G3BP1/2 in the formation of quiescent cells is through stress granule formation, the prediction is that wildtype and G3BP1/2 dKO U-2 OS cells should form a similar number of quiescent cells during sorbitol treatment. In contrast, if G3BP1/2 proteins promote cellular quiescence independently of stress granule, then wildtype cells are predicted to show higher levels of quiescence.

**Figure 4.**
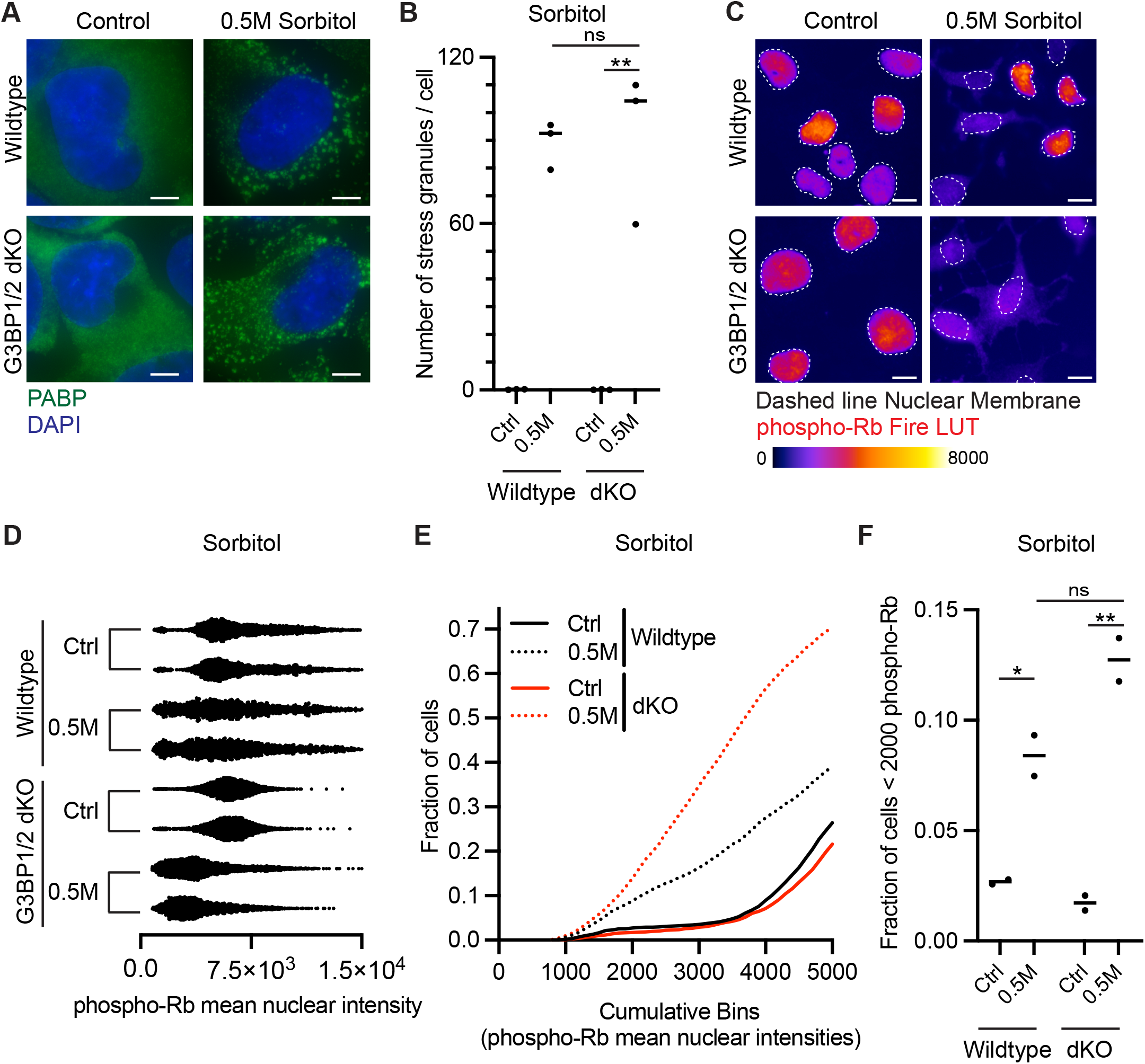
Sorbitol induces similar levels of low phospho-Rb cells between wildtype and G3BP1/2 dKO U-2 OS cells. **(A)** Representative PABP immunofluorescence images of wildtype and G3BP1/2 dKO U-2 OS treated with 0.5M sorbitol for 1 hour or left untreated. Scale bar = 5 µm. **(B)** Scatterplot of the number of stress granules per cell from wildtype and G3BP1/2 dKO U-2 OS cells treated with 0.5M sorbitol for 1 hour or left untreated. Three biological replicates were performed. 10 cells were counted for each biological replicate. ns = not significant; ** = P-value < 0.001. **(C)** Representative phospho-Rb immunofluorescence images of wildtype and G3BP1/2 dKO U-2 OS cells treated with 0.5M sorbitol for 6 hours or left untreated. Phospho-Rb immunofluorescence is converted to Fire LUT and the dashed lines indicate the nuclear membrane. Scale bar = 10 µm. **(D)** Mean phospho-Rb nuclear intensity scatterplot of wildtype and G3BP1/2 dKO U-2 OS from (C). Each dot represents a cell. At least 1742 cells were counted for each biological replicate. **(E)** Cumulative frequency plot of phospho-Rb mean nuclear intensities from 0 to 5000 phospho-Rb mean nuclear intensities from Figure 4D. **(F)** Scatterplot of fraction of cells with phospho-Rb mean nuclear intensity less than 5000 from Figure 4D. Two biological replicates were performed. ns = P-value > 0.05; * = P-value < 0.05; ** = P-value < 0.01.

Importantly, we observed a similar percentage of hypo-phosphorylated Rb cells between wildtype and G3BP1/2 dKO U-2 OS at 0.5M sorbitol (Figure 4D-E). These results argue stress granules, independent of G3BP1/2 genes, are involved in promoting cellular quiescence.

To extend this analysis, we examined using Hippuristanol to similarly trigger stress granule formation independently of G3BP1/2 proteins. Hippuristanol is an inhibitor of eIF4A, which can promote stress granules through two distinct mechanisms: Inhibiting translation (Bordeleau et al., 2006) and enhancing RNA-RNA interactions (Tauber *et al*., 2020). The latter mechanism can rescue stress granule formation in cells lacking G3BP1/2 proteins if 5 µM Hippuristanol is added to cells, but not at 1 µM, which inhibits translation but does not sufficiently inhibit eIF4A function in limiting intermolecular RNA-RNA interactions (Tauber *et al*., 2020). We discovered 5 µM, but not 1 µM, of Hippuristanol rescued stress granule formation in ΔΔG3BP1/2 U-2 OS cells (Figure 5A-B). Strikingly, we observed a similar percentage of cells with hypophosphorylated-Rb at 5 µM Hippuristanol between wildtype and G3BP1/2 dKO U-2 OS (Figure 5C-F). Conversely, 1 µM of Hippuristanol can robustly induce stress granule formation in wildtype U-2 OS cells, but not G3BP1/2 dKO U-2 OS cells (Figure 5A-B), and a greater fraction of hypophosphorylated-Rb cells are seen in wildtype U-2 OS cells compared to G3BP1/2 dKO U-2 OS cells (Figure 5C-F). Thus, restoration of stress granule formation in G3BP1/2 dKO U-2 OS cells by 5 µM Hippuristanol enhances the formation of quiescent cells providing additional evidence that stress granules are involved in promoting cellular quiescence.

**Figure 5.**
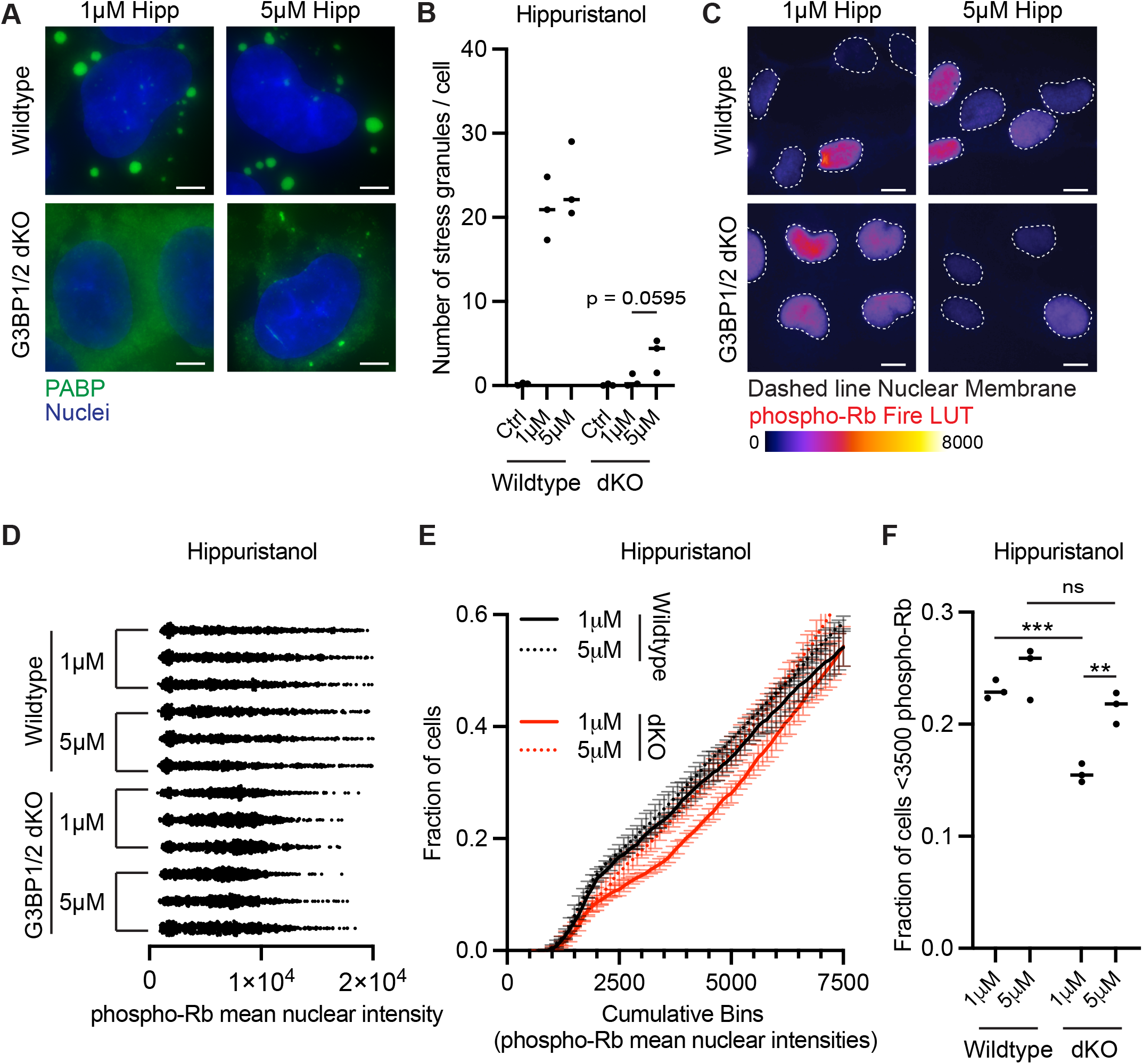
5 µM of Hippuristanol induce similar levels of low phospho-Rb between wildtype and G3BP1/2 dKO U-2 OS cells. **(A)** Representative PABP immunofluorescence images of wildtype and G3BP1/2 dKO U-2 OS cells when treated with 1 µM and 5 µM Hippuristanol for 1 hour. Scale bar = 5 µm. **(B)** Scatterplot of the number of stress granules per cell from wildtype and G3BP1/2 dKO U-2 OS cells treated with 1 µM and 5 µM Hippuristanol for 1 hour. Three biological replicates were performed. 10 cells were counted for each biological replicate. *** = P-value < 0.001. **(C)** Representative phospho-Rb immunofluorescence images of wildtype and G3BP1/2 dKO U-2 OS stained with antibodies binding to phospho-Rb protein when treated with 1 µM and 5 µM Hippuristanol for 6 hours. The phospho-Rb immunofluorescence is converted to Fire LUT and the dashed lines indicate the nuclear membrane. Scale bar = 10 µm. **(D)** Mean phospho-Rb nuclear intensity scatterplot of wildtype and G3BP1/2 dKO U-2 OS cells from Figure 5C. Each dot represents a cell. At least 684 cells were counted for each biological replicate. **(E)** Cumulative frequency plot of phospho-Rb mean nuclear intensities from 0 to 7500 from Figure 5D. Error bars represent the three biological replicates from Figure 5D. **(F)** Scatterplot of the fraction of cells with phospho-Rb mean nuclear intensity less than 3000 from Figure 5D. Three biological replicates were performed. ns = P-value > 0.05; *** = P-value < 0.001.

A prediction of the above experiment is that cells with increased numbers of stress granules would be more likely to be quiescent. We, and others (Majerciak et al., 2021), have observed that siRNA depletion, or CRISPR knockout of DDX6, leads to enhanced numbers of stress granules, and the formation of those stress granules is still dependent on G3BP1/2 (Figure 6A).

**Figure 6.**
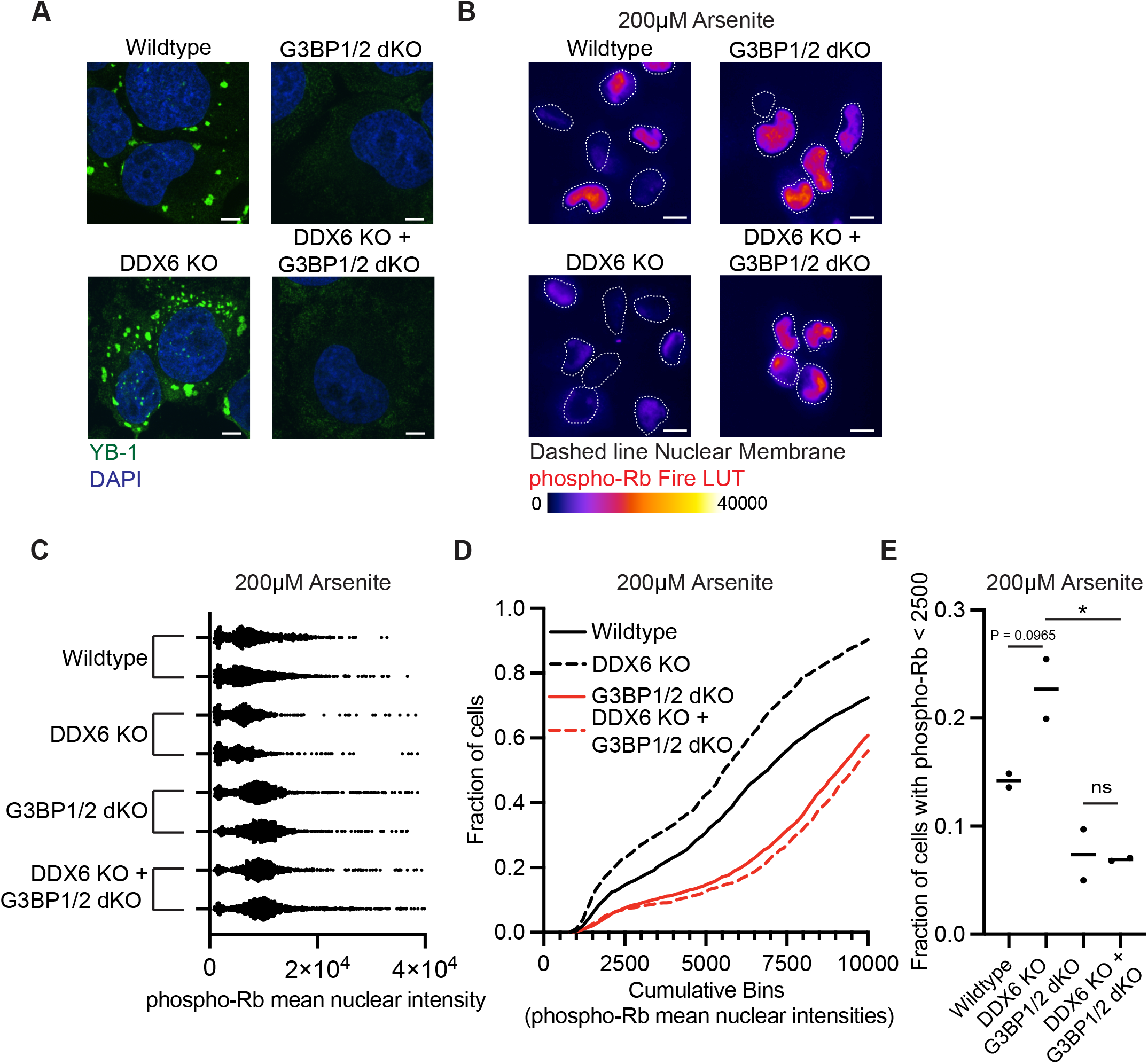
Knockout of DDX6 gene greater fraction of cells with low phospho-Rb under non-stress and arsenite conditions. **(A)** Representative YB-1 immunofluorescence images of wildtype, DDX6 KO, G3BP1/2 dKO, DDX6 KO + G3BP1/2 dKO U-2 OS cells when treated with 200 µM arsenite for 6 hours. The phospho-Rb immunofluorescence is converted to Fire LUT and the dashed lines indicate the nuclear membrane. Scale bar = 5 µm. **(B)** Representative phospho-Rb immunofluorescence images of wildtype, DDX6 KO, G3BP1/2 dKO, DDX6 KO + G3BP1/2 dKO U-2 OS cells when treated with 200 µM arsenite for 6 hours. The phospho-Rb immunofluorescence is converted to Fire LUT and the dashed lines indicate the nuclear membrane. **(C)** Mean phospho-Rb nuclear intensity scatterplot of wildtype, DDX6 KO, G3BP1/2 dKO, DDX6 KO + G3BP1/2 dKO U-2 OS cells when treated with 200µM arsenite. At least 811 cells were counted for each biological replicate. Scale bar **=** 10 µm **(D)** Cumulative frequency plot of phospho-Rb mean nuclear intensities from 0 to 10000 from Figure 6E. **(E)** Scatterplot of the fraction of cells with phospho-Rb mean nuclear intensity less than 2500 from Figure 6C. Two biological replicates were performed. ns = P-value > 0.05; * = P-value < 0.05.

Strikingly, we observed greater levels of stress granule formation and cells with hypophosphorylated-Rb in DDX6 KO U-2 OS cells compared to wildtype U-2 OS cells under arsenite conditions (Figure 6B-E). In addition, the increase in hypophosphorylated Rb in DDX6 KO U-2 OS is dependent on the stress granule G3BP1/2 genes (Figure 6A-E), which are required for the increased levels of stress granules seen in the DDX6 strain. These results provide additional evidence that stress granules promote cellular quiescence.

### Stress granules promote cellular quiescence in response to chemotherapeutics

Cellular quiescence (G0) is an important mechanism that facilitates the development of chemoresistance (Min and Spencer, 2019; Recasens and Munoz, 2019). Given this precedent, we examined how stress granules affected chemoresistance to vinorelbine, which is known to induce stress granules (Szaflarski *et al*., 2016). We asked if stress granules promote quiescence in the context of Vinorelbine by treating wildtype and G3BP1/2 dKO U-2 OS cells with 100 µM Vinorelbine for 6 hours. This concentration induced stress granules robustly in wildtype but not in G3BP1/2 dKO U-2 OS cells after 1 hour (Figure 7A-B) (Szaflarski *et al*., 2016). More importantly, we see fewer cells in quiescence in G3BP1/2 dKO U-2 OS cells compared to wildtype cells (Figure 7C-F). These results suggest stress granules promote cellular quiescence in the context of chemotherapeutics.

**Figure 7.**
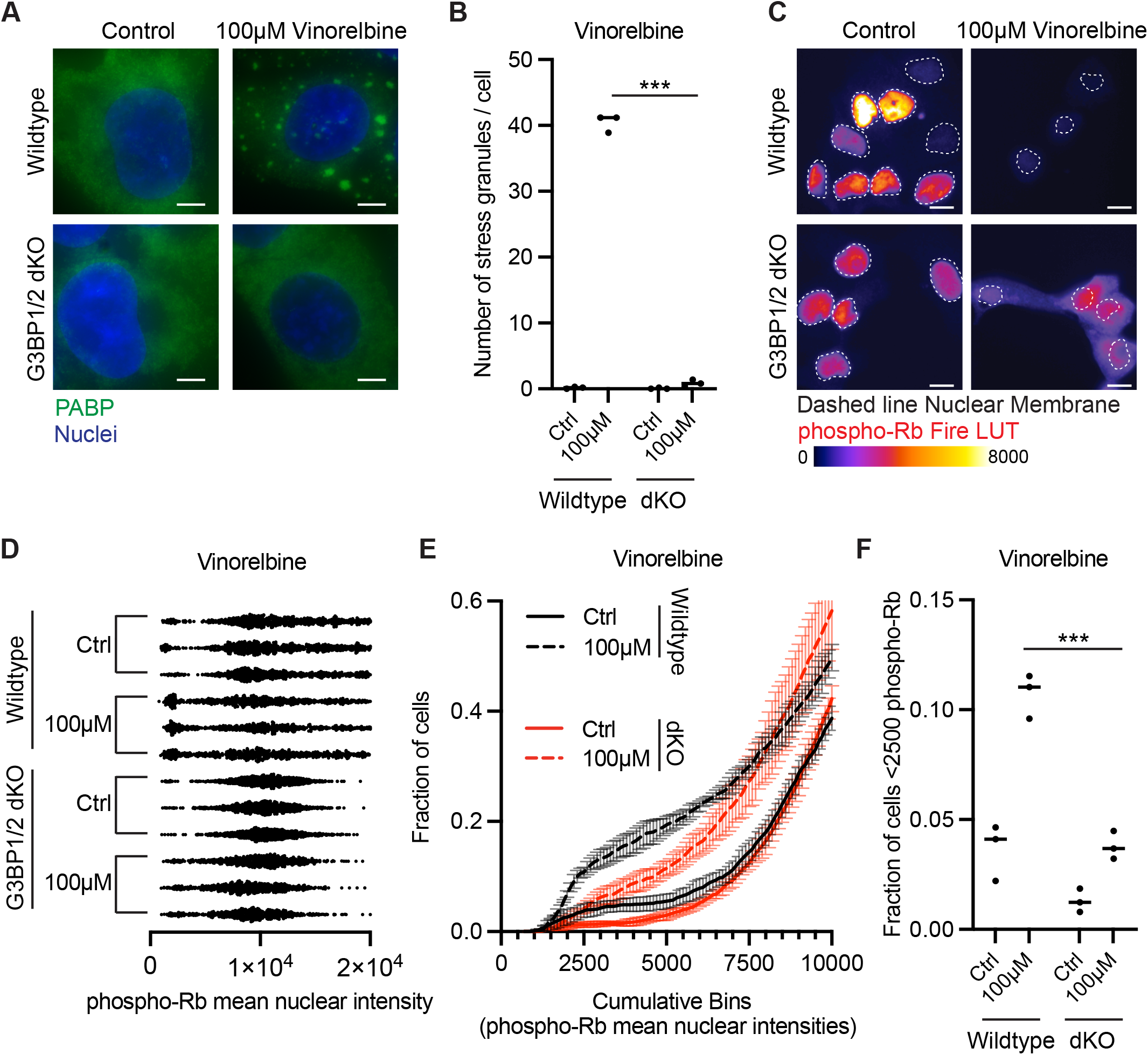
G3BP1/2 genes promote a greater fraction of cells with low phospho-Rb levels and greater levels of chemoresistance when treated with 100 µM Vinorelbine. **(A)** Representative PABP immunofluorescence images of wildtype and G3BP1/2 dKO U-2 OS cells when treated with 100 µM Vinorelbine for 1 hour. Scale bar = 5 µm. **(B)** Scatterplot of the number of stress granules per cell from wildtype and G3BP1/2 dKO U-2 OS cells treated with 100 µM Vinorelbine for 1 hour. Three biological replicates were performed. 10 cells were counted for each biological replicate. *** = P-value < 0.001. **(C)** Representative phospho-Rb immunofluorescence images of wildtype and G3BP1/2 dKO U-2OS cells when treated with 100 µM Vinorelbine for 6h. The phospho-Rb immunofluorescence is converted to Fire LUT and the dashed lines indicate the nuclear membrane. Scale bar **=** 10 µm **(D)** Mean phospho-Rb nuclear intensity scatterplot of wildtype and G3BP1/2 dKO U-2 OS cells. Each dot represents a cell. At least 622 cells were counted for biological replicate. **(E)** Cumulative frequency plot of phospho-Rb mean nuclear intensities from 0 to 10000 from Figure 6D. **(F)** Scatterplot of the fraction of cells with phospho-Rb mean nuclear intensity less than 2500 from Figure 7E. Two biological replicates were performed. *** = P-value < 0.001.

## DISCUSSION

In this manuscript, we provide several observations arguing stress granules are involved in promoting cellular quiescence. First, we demonstrate that cells with spontaneous stress granules that form during non-stress conditions are often quiescent. Second, we showed cells deficient in key stress granule assembly proteins (G3BP1/2) contain fewer quiescent cells under both stress and arsenite conditions. Third, we showed that this phenotype is dependent on stress granules because rescuing stress granule formation with 0.5M sorbitol or 5µM Hippuristanol in the G3BP1/2 dKO U-2 OS restores cellular quiescence levels to similar levels as wildtype U-2 OS cells. Fourth, we provided evidence that increasing stress granule assembly using a DDX6 KO U-2 OS cell line results in greater levels of cellular quiescence. We interpret these results to demonstrate that stress granule promotes entry into the quiescent cellular state.

Several observations argue that the ability of stress granules to enhance the formation of quiescent cells is one mechanism by which stress granules promote chemoresistance. First, Vinorelbine, a chemotherapeutic, induces a greater fraction of quiescent cells in wildtype U-2 OS cells versus G3BP1/2 dKO U-2 OS cells (Figure 7). A role for the quiescent state in promoting chemoresistance is consistent with many other works (Moore and Lyle, 2011; Recasens and Munoz, 2019). Since the quiescent state promotes resistance to many chemotherapeutic agents (Brown et al., 2017; Chen et al., 2012; Dembinski and Krauss, 2009; Min and Spencer, 2019; Naumov et al., 2003), and stress granules formation is triggered by many different types of chemotherapeutics including alkylating agents, anti-metabolites, mitotic inhibitors, kinase inhibitors, and proteasome inhibitors (Adjibade *et al*., 2015; Bittencourt *et al*., 2019; Fournier *et al*., 2010; Grabocka and Bar-Sagi, 2016; Kaehler *et al*., 2014; Lin *et al*., 2019; Szaflarski *et al*., 2016; Zhao *et al*., 2021), we suggest that the formation of quiescent cells by stress granule formation could be a general mechanism by which stress granules confer chemoresistance.

Determining if stress granules are involved in cell death can be difficult because inhibition of stress granules may also perturb other unrelated stress granule processes. For example, one type of experimental evidence that is used to argue stress granules play a role in cell death is by inhibiting stress-induced translational shutoff (Adjibade et al., 2020; Adjibade *et al*., 2015; Fournier *et al*., 2010; Szaflarski *et al*., 2016). However, this evidence is not particularly convincing because it does not distinguish translational shutoff versus stress granule assembly. Alternatively, knockdown or knockout of key stress granule regulators that sensitize cells to chemotherapeutic-induced cell death can also be problematic (Fournier *et al*., 2010; Kaehler *et al*., 2014; Lin *et al*., 2019; Szaflarski *et al*., 2016). For example, G3BP1/2 proteins have roles independent of stress granule assembly including enhancing translation of short RNAs, rescuing stalled 40S ribosomes, and inhibiting mTORC1 signaling (Laver *et al*., 2020; Meyer *et al*., 2020; Prentzell *et al*., 2021). It remains possible that G3BP1/2 proteins, and/or stress granules will affect chemoresistance by other mechanisms, although our data argues strongly that promoting quiescence is one mechanism of stress granule induced chemoresistance.

Stress granules have been proposed to limit cell death by other mechanisms. These include sequestration of pro-apoptotic proteins and reduction of reactive oxygen species (Amen and Kaganovich, 2021; Arimoto *et al*., 2008; Kim *et al*., 2005; Takahashi *et al*., 2013; Thedieck *et al*., 2013; Tsai and Wei, 2010; Wippich *et al*., 2013; Zhao *et al*., 2021). However, it is unclear if many of these proposed mechanisms are restricted to certain stressors and chemotherapeutics or if they can be generalized. In fact, under Vinorelbine treatment, unlike arsenite, several proapoptotic factors are not sequestered to stress granules (Szaflarski *et al*., 2016). Therefore, we suggest stress granule-induced cellular quiescence can be a potential explanation for cell survival for many stressors and chemotherapeutics.

One major implication from this work is it may explain the numerous multiple mechanisms by which stress granules limit apoptosis. Both stress granules and cellular quiescence can limit reactive oxygen species levels during cellular stress (Amen and Kaganovich, 2021; Marescal and Cheeseman, 2020; Takahashi *et al*., 2013). Therefore, some of the reduction of reactive oxygen species levels attributed to stress granules could be through stress granule induction of cellular quiescence. In addition, quiescent cells are transcriptionally reprogramed to downregulate genes that promote apoptosis (Coller *et al*., 2006). Therefore, it is possible some of the mechanisms proposed for how stress granules regulate apoptosis are actually downstream consequences of the induction of cellular quiescence. Clearly, further work is required to disentangle how stress-granule induced quiescence is altering/limiting apoptosis which may provide a general explanation for how stress granules reduce cell death.

The ability of cancer cells to enter the quiescence state can negatively affect disease outcomes. This includes chemotherapeutic resistance (Chen et al., 2016; Recasens and Munoz, 2019). In addition, it has also been recognized that quiescent tumor cells can be a source of metastasis. Dormant cancer cells can metastasis to distant areas, remain quiescent, and reenter the cell cycle for up to 20 years (Goddard et al., 2018). G3BP1/2 genes were shown recently to be involved in metastasis (Somasekharan *et al*., 2015) which may provide an intriguing link between stress granules, cellular quiescence, and metastasis. Recognizing the importance of quiescence in tumors, recent therapies are being developed to “Lock out” cancer cells from quiescence which renders the tumors susceptible to chemotherapeutics (Recasens and Munoz, 2019). Our model suggests targeting stress granules may provide another therapeutic strategy to “Lock out” cancer cells from cellular quiescence.

## ACKNOWLEDGEMENTS

We would like to thank Paul Anderson’s lab (Brigham Children’s Hospital) for providing us with wildtype and ΔΔG3BP1/2 U-2 OS cell lines. We like to thank Carolyn Decker for DeltaVision training and Joe Dragavon (BioFrontiers Institute Advanced Light Microscopy Core; RRID: SCR_018302) for PerkinElmer Opera Phenix and Spinning Disk Confocal Nikon Ti-E microscope training and image analysis help. We would also like to thank Theresa Nahreini for training and help with cell culture (Cell Culture Facility). This work was funded by Howard Hughes Medical Institute (R.P.) and a Banting Postdoctoral Fellowship (A.K.).

## AUTHOR CONTRIBUTIONS

A.K., N.R., and L.M.V. conceived and performed experiments and analyzed results. A.K., S.S., and R.P. contributed to manuscript preparation. R.P. contributed to project conception.

## MATERIALS AND METHODS

### Cell lines

Wildtype and G3BP1/2 dKO U-2 OS are kindly provided to us by Paul Anderson’s lab at Brigham Children’s Hospital (Kedersha *et al*., 2016). GFP-G3BP1 + G3BP1/2 dKO U-2 OS cell-line was generated in the Parker lab using a lentivirus expressing GFP-G3BP1 that was established by James Burke (Burke et al., 2020). DDX6 KO U-2 OS and DDX6 KO + G3BP1/2 dKO U-2 OS cell lines were also generated in the Parker Lab (Ripin et al., 2022).

### Cell’s growth conditions

U-2 OS cells were maintained in DMEM with 10% FBS and 1% Penicillin/Streptomycin at 37°C/5% CO_2_.

### Generation of DDX6 knockout cell lines

The CRISPR/Cas9 guide RNAs targeting two regions within the DDX6 locus were designed using the Integrated DNA Technologies (IDT) CRISPR guide target design tool. Overlapping oligos (DDX6 sgRNA 1, 2 sense and DDX6 sgRNA 1,2 antisense (table with sequences) were annealed in T4 DNA ligase buffer and ligated into the Bbs1 sites in pSpCas9(BB)-2A-GFP (px458) using T4 DNA ligase.

To generate DDX6 knockout in U-2 OS and G3BP1/2 dKO U-2 OS lines, cells (T-25 flask; 60% confluent) were co-transfected with 3 µg px458-DDX6 and 400-ng of pcDNA3.1-puro using 15-µl of Lipofectamine 2000 (Thermo Fisher Scientific) according to manufacturer’s instructions. 24 hours post-transfection, Cas9-GFP expression was observed via fluorescent microscopy. The medium was replaced with a medium containing 2 µg/ml of puromycin. Selective medium was replaced 2 days post-transfection. Five days post-transfection, selective growth medium was replaced with normal growth medium. When cells reached 80% confluency, cells were serial diluted and plated on 15-cm dishes. Individual colonies were isolated, propagated, and screened via immunoblot analysis.

### Immunofluorescence

#### (A) Analysis of individual phospho-Rb nuclear intensities

10,000 U-2 OS cells were seeded on CellCarrier-96 Ultra Microplates and incubated overnight. Cells were either untreated or treated with various compounds for 6 hours as noted in each figure. Cells were then washed once with PBS, fixed with 4% PFA for 5 mins, and permeabilized with 0.1% Triton X-100 in PBS for 5 mins. Cells were stained and incubated with primary antibody 1:250 rabbit anti-phospho-Rb (Cell signaling; 8516) for 1 hour. Cells were then washed 3x with PBS. Subsequently, cells were stained with goat anti-rabbit Alexa Fluor 657 (Abcam; ab150079)). Cells were washed 3x with PBS. Stained for 5 mins with 1:2000 DAPI (Thermo Fisher Scientific; 62248) in PBS. Wash 2x with PBS and store in PBS at 4C.

#### (B) Analysis of stress granules

200,000 U-2 OS cells were seeded in 6-well plates containing EtOH-treated coverslips. Cells were either untreated or treated with various compounds for 1 hour as noted in each figure. Cells were then washed once with PBS, fixed with 4% PFA for 5 mins, and permeabilized with 0.1% Triton X-100 in PBS for 5 mins. Cells were stained and incubated with primary antibodies (1:1000 rabbit anti-PABP (Abcam; ab21060) or 1:500 rabbit anti-YBX1 (Proteintech; 20339-1-AP) for 1 hour. Cells were then washed 3x with PBS. Subsequently, cells were stained with secondaries (goat anti-rabbit FITC (Abcam; ab6717) or goat anti-rabbit Alexa Fluor 647 (Abcam; ab150079)). Cells were washed 3x with PBS. Stained for 5 mins with 1:2000 DAPI (Thermo Fisher Scientific; 62248) in PBS. Wash 2x with PBS and mounted on slides with Vectashield Antifade Mounting Medium (Vector Laboratories, H-1000).

### Imaging

#### (A) Analysis of individual phospho-Rb nuclear intensities

Cells were imaged with the PerkinElmer Opera Phenix at the BioFrontiers Advanced Light Microscopy Core. Cells were imaged using the 40X 1.1-NA water objective (PerkinElmer; HH1400422). Appropriate lasers, transmission, and exposure times were used to best capture images and were kept consistent within each experiment. Images are exported as tiff files. All images shown are single z-slice and altered with Fire LUT with the help of ImageJ with FIJI plugin and Adobe Photoshop.

#### (B) Analysis of stress granules

Cells were imaged using a GE widefield DeltaVision Elite microscope with an Olympus UPlan-SApo 100X 1.40-NA objective Oil Objective lens and a PCO Edge sCMOS camera with the help of SoftWoRx Imaging software or a Spinning Disk Confocal Nikon Ti-E microscope with a 100X 1.45-NA objective lens and a 2x Andor Ultra 8888 EMCCD camera. Appropriate lasers, transmission, exposure times were used to best capture images and were kept consistent within each experiment. All images shown are z-stacked images of entire cells using ImageJ with FIJI plugin and Adobe Photoshop.

### Image analysis

#### (A) Analysis of individual phospho-Rb nuclear mean intensities

Exported tiff files were imported into Imaris 9.8.2 version for image analysis. Nuclear mean intensity for phospho-Rb was quantified using the surface tool. Briefly, the surface tool was used to define the nucleus by the DAPI stain. The tool was applied identically for every compared sample. Nuclear mean intensities from phospho-Rb and p21 channels were then collected from the surfaces. In Figure 6, phospho-Rb nuclear mean intensities was performed using MATLAB and ImageJ with Fiji plug-in custom batch scripts.

#### (B) Analysis of stress granules

Deltavision files were imported directly into Imaris 9.8.2 version for image analysis. Stress granules were identified and quantified using the surface tool with PABP fluorescence. The tool was applied identically for every compared sample. Stress granule positive cells from Figure 1 were determined manually by visual inspection.

